# Transcranial alternating current stimulation reverses adolescent alcohol-induced cellular and behavioral deficits

**DOI:** 10.64898/2026.10.30.763090

**Authors:** E.D.K. Sullivan, A. Bresnicky, H.B. Trevathan, L.F. Gallagher, R. Solanki, F. Fröhlich, D.L. Robinson

## Abstract

**BACKGROUND:** Alcohol is commonly consumed by adolescents in a binge-like pattern that can produce persistent cognitive impairments. Previous work found that adolescent intermittent ethanol (AIE) exposure reduces parvalbumin expressing (PV+) interneurons and increases perineuronal net (PNN) expression in brain regions critical for cognition. Transcranial alternating current stimulation (tACS), a noninvasive neuromodulation technique, has shown promise for treating psychiatric disorders associated with cognitive dysfunction. Here, we examined whether AIE exposure alters PV+ interneurons and PNN expression in the medial prefrontal cortex (mPFC) and impairs cognition, and whether mPFC-targeted gamma tACS can reverse these effects.

**METHODS:** Sprague Dawley rats (N=139, 50% female) underwent AIE or control exposure during adolescence and received Active or Sham mPFC-targeted gamma tACS treatment for three consecutive days in adulthood. Following stimulation, rats either underwent immunohistochemical assessment of PV+ and PNN expression or behavioral testing using the novel object recognition (NOR) task.

**RESULTS:** AIE exposure reduced PV+ cell counts in the prelimbic cortex, an effect that was reversed by active tACS. In contrast, AIE exposure did not alter PNN measures. Interestingly, active tACS affected the prelimbic and infralimbic cortices more than the anterior cingulate cortex. AIE exposure also reduced recognition memory of a novel object, and this deficit was ameliorated by active tACS when assessed one week after stimulation. Additionally, active stimulation decreased anxiety-like behavior in the open-field test.

**CONCLUSIONS:** Together, these results confirm that adolescent alcohol exposure produces lasting cellular and cognitive alterations and supports mPFC-targeted gamma tACS as a potential intervention for alcohol-induced dysfunction.

## INTRODUCTION

Alcohol is one of the most widely used drugs by adolescents (1) and is often consumed in a binge-drinking pattern (2). Binge drinking during adolescence is particularly concerning because this developmental period is characterized by extensive neurobiological maturation, rendering the brain especially vulnerable to insults (3). Importantly, adolescent binge drinking can produce persistent impairments in neurocognitive function (2), for which there are few effective treatments.

A substantial body of preclinical research has demonstrated that adolescent intermittent ethanol (AIE) exposure produces long-term behavioral impairments in adulthood, including increased anxiety-like behavior, deficits in object and spatial memory, impaired behavioral flexibility, and disrupted social interactions (4–10). Although the neurocircuitry underlying these outcomes is complex, the medial prefrontal cortex (mPFC) is critical in cognitive processes such as learning, memory, and executive function. As the mPFC continues to mature throughout adolescence, it is particularly susceptible to alcohol-induced neurodevelopmental disruptions (11,12).

Parvalbumin-positive (PV+) interneurons are fast-spiking inhibitory neurons that play a central role in regulating cortical network activity and generating gamma-frequency oscillations (13–15). Gamma oscillations are integral for information processing and contribute to numerous higher-order cognitive functions, including memory (16,17). Notably, many cortical PV+ neurons are ensheathed by perineuronal nets (PNNs) (18,19), specialized extracellular matrix structures that stabilize synaptic connections (16) and protect neurons from oxidative stress (20). Consistent with a role for PV+ interneurons and PNNs in alcohol-induced cognitive dysfunction, we previously reported that AIE exposure reduced PV+ cell size within the anterior insular cortex while increasing the proportion of PV+ neurons surrounded by PNNs. Mediation analyses further indicated that the ratio of PV+ cells surrounded by PNNs and the number of PNNs statistically contributed to the AIE-induced behavioral deficits (21). These findings suggest that long-lasting alterations in PV+ interneurons and their PNN regulation may represent a mechanistic link between adolescent alcohol exposure and persistent cognitive impairments, highlighting this cellular population as a potential therapeutic target.

One intervention with the potential to reverse AIE-induced disruptions is transcranial alternating current stimulation (tACS), a noninvasive neuromodulation technique that delivers low-amplitude alternating electrical current through the scalp to modulate neural activity. A key advantage of tACS is its ability to be administered at frequencies corresponding to endogenous brain rhythms, such as alpha, theta, or gamma frequencies (22), which play critical roles in cognitive function. For example, alpha oscillations are associated with perception, while theta and gamma rhythms contribute to cognitive control. Notably, disrupted gamma activity has been identified in several psychiatric disorders (23). Clinically, tACS has shown therapeutic potential for schizophrenia (24), depression (25,26), and addiction (27,28), with addiction history moderating the effect of prefrontal tACS administration (29). In particular, gamma-frequency tACS has been reported to improve mood and cognitive performance in patients with major depressive disorder (30) and shows promise for improving episodic memory and cognitive performance in those with Alzheimer’s disease or mild cognitive impairment (31–34), although these effects appear to be dependent on stimulation parameters.

The therapeutic effects of tACS are thought to arise, at least in part, from the entrainment and re-synchronization of neural oscillations that are disrupted in psychiatric disorders (30). At the cellular level, PV+ interneurons may be particularly sensitive to tACS (35,36). For instance, tACS entrained alpha oscillations by synchronizing fast-spiking cortical neurons in ferrets (36), and 80-Hz gamma tACS restored coherence measures between the mPFC and nucleus accumbens in rats that had cocaine-induced hypoactivity (37). Given the central role of PV+ interneurons in generating and maintaining gamma oscillations, these findings raise the possibility that gamma-frequency tACS may ameliorate alcohol-induced cognitive deficits by altering PV+ interneurons and their surrounding PNNs.

Therefore, we hypothesized that adolescent alcohol exposure produces long-lasting alterations in PV+ interneurons and their associated PNNs while impairing cognition. Furthermore, we hypothesized that gamma-frequency tACS would alter PV+ interneurons and improve AIE-induced memory deficits. To test these hypotheses, we quantified the number and size of PV+ interneurons and associated PNNs within the anterior cingulate (ACC), prelimbic (PL), and infralimbic (IL) cortices of rats following AIE exposure and subsequent mPFC-targeted gamma tACS treatment in adulthood. Next, we assessed the effects of AIE exposure and gamma tACS on cognition using the novel object recognition (NOR) task in rats.

## METHODS AND MATERIALS

### Subjects and alcohol exposure paradigm

Male and female Sprague-Dawley rats (N=139) were bred in-house, weaned on postnatal day (PD) 21, and housed in same-sex littermate pairs. Rats were maintained on a 12-hour light/dark cycle (lights on at 0700) with unrestricted access to food and water, except as described below (see “Transcranial alternating current stimulation”). All rats received either the adolescent intermittent ethanol (AIE) paradigm (5 g/kg of 25% w/v ethanol in water) or a volume-matched water control (CON) via oral gavage on a 2-days-ON/2-days-OFF schedule from PD25 to PD54. We previously showed that this AIE schedule produces blood alcohol concentrations of ∼230 mg/dL in PD35 rats (38). All procedures were approved by the Institutional Animal Care and Use Committee at the University of North Carolina at Chapel Hill and complied with National Institute of Health regulations for the care and use of laboratory animals.

### Experimental design

These studies used a 2 (Exposure; AIE, CON) X 2 (tACS Treatment; Active, Sham) design such that four groups were compared in each of the three experiments: CON+Sham, CON+Active, AIE+Sham, AIE+Active (Figure 1A). Group sizes ranged from 9 to 14 rats per group in each experiment and were balanced across sex. Rats in Experiment 1 (N=48) were perfused 90 minutes after the third stimulation session. Rats in Experiment 2 (N=53) received stimulation in the morning and underwent Novel Object Recognition (NOR) testing that afternoon. Rats in Experiment 3 (N=38) underwent NOR testing one week after the final stimulation session.

**Figure 1.**
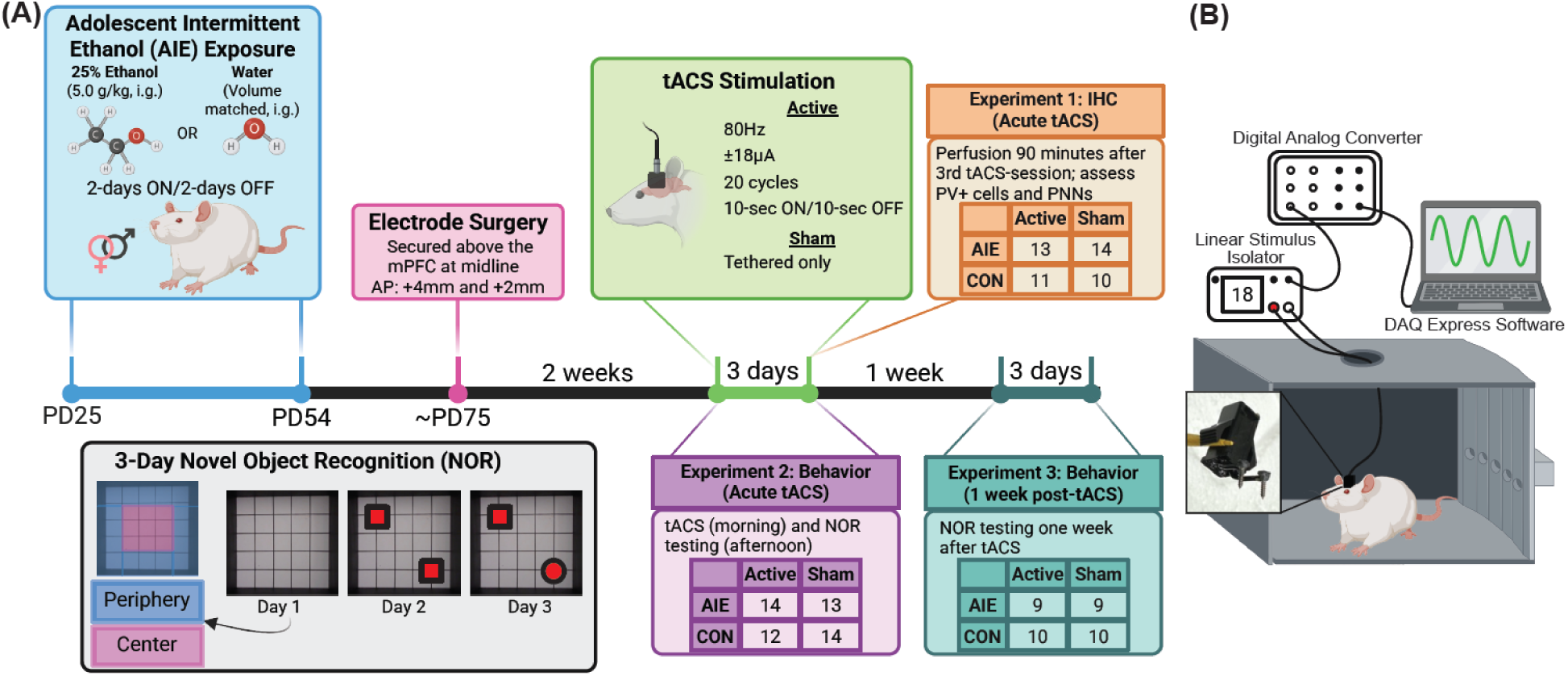
Experimental timeline. (A) Male and female rats underwent AIE exposure or CON gavage from postnatal day (PD) 25-54. Once adult, the tACS electrode was surgically attached, and rats were allowed to recover for 2 weeks. Active tACS stimulation or Sham stimulation occurred for three consecutive days. Animals in Experiment 1 were perfused 90 minutes following the third stimulation session for immunohistochemistry (IHC). Animals in Experiment 2 underwent novel object recognition (NOR) testing on the same days of stimulation. Animals in Experiment 3 underwent NOR one week post-stimulation. Rat numbers (approximately 50% female) are shown for each experiment in the corresponding table. (B) Diagram of the tACS setup. DAQ Express Software generated the waveform which passed through a digital analog converter and linear stimulus isolator. The animals’ custom-built electrodes were tethered to the linear stimulus isolator. Sham animals were tethered for the same amount of time as active tACS rats, but no current was passed. Created in BioRender. Sullivan, E. (2026) https://BioRender.com/n0idq58.

### Electrode preparation and stereotaxic surgery

tACS electrodes were designed and constructed in-house, and each electrode’s electrical continuity was verified using an oscilloscope prior to being implanted. About PD75, animals were anesthetized with isoflurane (4% induction, 2.5% maintenance) and secured in a stereotaxic frame. The skull was exposed and cleaned. The tACS electrodes consisted of two stainless-steel screws (Fine Science Tools, Foster City, CA) placed on the surface of the skull (i.e., non-penetrating) along midline (from bregma: AP +2mm and +4mm) to target the mPFC. The tACS electrode assembly was secured with additional skull screws and dental cement.

### Transcranial alternating current stimulation

Following a one-week post-surgery recovery period, rats were mildly food restricted to maintain weight for the remainder of the experiment. This was implemented to match our previously published studies in which food restriction was necessary for appetitive behavior paradigms (6,8,21).

Prior to stimulation, rats were habituated to the tether and stimulation chamber (Med Associates, St. Albans, VT) with the white noise, house light, and chamber fan activated. Stimulation sessions occurred in the morning across three consecutive days. Rats assigned to active tACS received an 80 Hz sinusoidal waveform consisting of 20 cycles of 10-seconds-ON/10-seconds-OFF at 18 µA (Figure 1B) (37). Rats assigned to sham tACS underwent identical handling and tethering, but no current was delivered. In a subset of animals, videos were acquired during the first active or sham stimulation session and line crossings were quantified by an experimenter blinded to condition.

### Tissue collection and preparation

Ninety minutes after the third stimulation session, rats in Experiment 1 were anesthetized with isoflurane and transcardially perfused with 0.1M phosphate-buffered saline (1X PBS) followed by 4% paraformaldehyde (PFA) in 1X PBS. Brains were extracted and post-fixed in 4% PFA at 4°C for at least 24 hours before being transferred to 30% sucrose in 1X PBS for a minimum of 3 days. The brains were then frozen on dry ice and coronally sectioned in 40-µm sections using a microtome (HM 450, ThermoScientific). Slices were collected sequentially into 12-well plates and stored at -20°C in cryoprotectant solution (30% glucose, 30% ethylene glycol in 1X PBS) until further processing.

### Immunofluorescence

Prefrontal cortical slices (+4.20 to +2.52 mm from bregma) (39) were selected to assess PV+ neurons and PNNs in the ACC, PL, and IL. For each rat in Experiment 1, 4-6 representative sections per region were analyzed, allowing for systematic sampling along the anterior-posterior axis (40). Slices were transferred to reagent trays and washed five times for 5-minutes each in 1X PBS to remove cryoprotectant. Slices were then placed in a 5% blocking solution (200 µL Triton X-100, 5 mL normal Goat serum, 95 mL 1X PBS) for 1 hour at room temperature. Next, slices were incubated for 24 hours at 4°C on an orbital shaker in a primary antibody solution containing rabbit anti-PV (1:500 dilution; Abcam Cat. #AB11427) prepared in 5% blocking solution. Following three 5-minute washes in 1X PBS, slices were incubated under foil for 2 hours in a secondary antibody solution containing goat anti-rabbit Alexa Fluor 594 (1:1000 dilution; Thermo Fisher Cat. #A32740) and WFA lectin fluorescein (1:500 dilution; Vector Laboratories, SKU FL-1351-2) prepared in 5% blocking solution. After three additional 5-minute washes in 1X PBS, slices were mounted and coverslipped with ProLong Gold Antifade Mountant (Thermo Fisher Cat. #P36960).

Slices were visualized using a DS-RiZ scope (Nikon, Inc.) and quantified with NIS-Elements AR46 (Nikon, Inc.) following previously described methods (21,40). Images were acquired as z-stacks containing 7 steps (2.5 µm/step) with the ACC, PL, and IL captured in a single image.

PV+ neurons and PNNs were quantified as total object number normalized to area and mean object size. Co-labeled immunofluorescence was used to determine the proportion of PV+ neurons ensheathed by PNNs and the proportion of PNNs containing PV+ neurons.

### Experimenters were blinded to group conditions

#### Novel object recognition task

All rats in Experiments 2 and 3 underwent a three-day NOR paradigm performed in the afternoon on either the stimulation day (Experiment 2) or one week after stimulation (Experiment 3). Behavioral testing was conducted in an open-field arena (66 x 66 cm) marked with a grid of 25 equally-sized boxes on the floor and was in a room separate from the housing room. Room lighting was dimmed and maintained at approximately 40 lux. A video camera was used to record each session. At the start of each testing day, rats were habituated to the testing room for 15 minutes while remaining in their home cages. On Day 1, rats were placed in the center of the arena and allowed to freely explore for 10 minutes. On Day 2, two identical novel objects were placed in opposite corners of the arena, and rats were again allowed to freely explore for 10 minutes. On Day 3, one familiar object was replaced with a novel object, and rats explored the arena for 10 minutes. Object exploration was manually scored and defined as touching, sniffing, biting, or climbing on an object. The arena was thoroughly cleaned between subjects. Novel object preference was calculated as: (Time spent exploring the novel object / Total object exploration time) X 100. Rats that spent less than 45 seconds exploring both objects combined on Day 2 were excluded from analysis.

#### Open-field analyses

Anxiety-like behavior was assessed on Day 1 of the NOR paradigm using latency to enter the center zone of the open-field arena and time spent in the center zone. The center zone was defined as the nine central squares of the arena. A center entry was defined as all four paws crossing into the center zone. Locomotor activity was assessed by the total number of line crossings. Latency to enter the center zone, time spent in the center zone, and number of line crossings were manually scored by an experimenter blinded to condition.

#### Statistical analyses

For Experiment 1, generalized linear models (GLMs) were applied to the immunofluorescent data with Exposure (AIE, CON) and Treatment (Active, Sham) as factors and Sex (Male, Female) as a covariate. Outliers were identified as values greater than two standard deviations from the group mean. For Experiments 2 and 3, behavioral data were analyzed using two-way analyses of variance (ANOVAs) with Exposure (AIE, CON) and Treatment (Active, Sham) as between-subjects factors and Sex (Male, Female) as a covariate. Behavioral outliers were identified using Tukey’s method. All analyses were conducted in IBM SPSS Statistics (Version 32.0.0.0). Statistical significance was set at α = 0.05.

## RESULTS

### Locomotor activity during stimulation

We evaluated whether tACS altered spontaneous locomotion. There were no group differences (Active vs. Sham) in the number of line crossings during the first day of stimulation (data not shown).

### Immunofluorescence

We assessed whether three days of tACS altered PV+ or PNN immunofluorescence in three subregions of the mPFC directly below the electrode. Full statistics are shown in Table 1.

**Table 1.** All immunohistochemistry results. Generalized linear models were applied to the immunofluorescent data with Exposure (AIE, CON) and Treatment (Active, Sham) as factors and Sex (Male, Female) as a covariate. The Wald Chi-Square degree of freedom = 1 for each variable. Significant p-values (p < 0.05) are bolded and red.

| **Immunohistochemistry** |  |  |  |  |  |  |
| **a. Anterior Cingulate Cortex** |  |  |  |  |  |  |
| **Variable** | **Effect** | **Wald Chi-square** | ***p*** | **Exp(B)** | **95% Wald Confidence Interval** |  |
|  |  |  |  |  | **Lower** | **Higher** |
|---|---|---|---|---|---|---|
| PV+ Num/Area | Exposure | 1.426 | 0.232 | <0.001 | 6.79E-13 | 899.052 |
|  | Treatment | 5.879 | **0.015** | <0.001 | 2.96E-19 | 0.011 |
|  | Sex | 1.372 | 0.242 | <0.001 | 5.87E-14 | 2155.539 |
|  | Exp. X Treatment | 5.401 | **0.02** | 8.51E+12 | 106.078 | 6.83E+23 |
| PV+ Cell Size | Exposure | 3.372 | 0.066 | 2785694508 | 0.232 | 3.35E+19 |
|  | Treatment | 1.062 | 0.303 | 470484.929 | 7.66E-06 | 2.89E+16 |
|  | Sex | 0.709 | 0.4 | 43073.409 | 7.01E-07 | 2.65E+15 |
|  | Exp. X Treatment | 3.04 | 0.081 | <0.001 | 1.26E-28 | 42.737 |
| PNN Num/Area | Exposure | 0.017 | 0.895 | 0.213 | 2.30E-11 | 1961252357 |
|  | Treatment | 1.141 | 0.285 | <0.001 | 9.99E-17 | 51518.108 |
|  | Sex | 0.298 | 0.585 | 0.001 | 5.77E-14 | 29748573.56 |
|  | Exp. X Treatment | 2.262 | 0.133 | 50805490556 | 0.001 | 4.54E+24 |
| PNN Size | Exposure | 0.001 | 0.977 | 0.604 | 8.81E-16 | 4.14E+14 |
|  | Treatment | 0.123 | 0.726 | 0.001 | 6.90E-20 | 2.21E+13 |
|  | Sex | 0.535 | 0.465 | <0.001 | 4.85E-23 | 15476345722 |
|  | Exp. X Treatment | 0.153 | 0.696 | 18376.612 | 7.37E-18 | 4.58E+25 |
| Number of PV+ cells surrounded by PNNs | Exposure | 0.364 | 0.546 | 0.075 | 1.64E-05 | 340.388 |
|  | Treatment | 1.348 | 0.246 | 0.004 | 4.16E-07 | 43.009 |
|  | Sex | 0.045 | 0.831 | 0.367 | 3.61E-05 | 3731.106 |
|  | Exp. X Treatment | 1.376 | 0.241 | 1769.833 | 0.007 | 471946857.7 |
| PV+ cell size when surrounded by PNNs | Exposure | 2.295 | 0.13 | 1819906541 | 0.002 | 1.74E+21 |
|  | Treatment | 0.262 | 0.609 | 2229.642 | 3.36E-10 | 1.48E+16 |
|  | Sex | 0.372 | 0.542 | 9782.354 | 1.48E-09 | 6.49E+16 |
|  | Exp. X Treatment | 1.957 | 0.162 | <0.001 | 8.48E-31 | 105463.291 |
| Number of PNNs surrounding PV+ cells | Exposure | 0.399 | 0.528 | 0.072 | 2.11E-05 | 249.018 |
|  | Treatment | 1.14 | 0.286 | 0.008 | 1.04E-06 | 58.052 |
|  | Sex | 0.024 | 0.877 | 0.494 | 6.60E-05 | 3690.365 |
|  | Exp. X Treatment | 1.344 | 0.246 | 1265.121 | 0.007 | 222388055.6 |
| PNN size when surrounding PV+ cells | Exposure | 0.368 | 0.544 | 6645827.543 | 6.15E-16 | 7.19E+28 |
|  | Treatment | 0.046 | 0.831 | 0.003 | 6.96E-27 | 1.02E+21 |
|  | Sex | 0.042 | 0.838 | 0.003 | 9.04E-27 | 1.32E+21 |
|  | Exp. X Treatment | 0.155 | 0.694 | <0.001 | 2.61E-38 | 1.05E+25 |
| Ratio of PV+ cells surrounded by PNNs (PV+PNN/PV total) | Exposure | 0.015 | 0.903 | 0.569 | 6.72E-05 | 4819.462 |
|  | Treatment | 0.023 | 0.88 | 2.102 | 0 | 33586.452 |
|  | Sex | 0.888 | 0.346 | 105.028 | 0.007 | 1678543.077 |
|  | Exp. X Treatment | 1.668 | 0.197 | 4751.698 | 0.013 | 1805535362 |
| Ratio of PNNs that surround PV+ cells (PNN+PV/PNN total) | Exposure | 0.03 | 0.863 | 2.408 | 0 | 53652.781 |
|  | Treatment | 0.103 | 0.749 | 0.174 | 3.85E-06 | 7813.066 |
|  | Sex | 0.05 | 0.823 | 0.275 | 3.42E-06 | 22104.827 |
|  | Exp. X Treatment | 0.085 | 0.77 | 0.113 | 4.82E-08 | 263348.531 |
| **b. Prelimbic Cortex** |  |  |  |  |  |  |
| **Variable** | **Effect** | **Wald Chi-square** | ***p*** | **Exp(B)** | **95% Wald Confidence Interval** |  |
|  |  |  |  |  | **Lower** | **Higher** |
| PV+ Num/Area | Exposure | 2.209 | 0.137 | <0.001 | 3.08E-20 | 481.259 |
|  | Treatment | 2.483 | 0.115 | <0.001 | 3.77E-22 | 213.332 |
|  | Sex | 0.097 | 0.755 | 0.013 | 1.71E-14 | 9671321109 |
|  | Exp. X Treatment | 4.607 | **0.032** | 1.84E+17 | 31.611 | 1.07E+33 |
| PV+ Cell Size | Exposure | 0.603 | 0.437 | 2535.197 | 6.54E-06 | 9.82E+11 |
|  | Treatment | 0.067 | 0.795 | 0.052 | 1.06E-11 | 253831663.4 |
|  | Sex | 0.218 | 0.641 | 0.006 | 4.17E-12 | 10073334.33 |
|  | Exp. X Treatment | 0.114 | 0.736 | 0.007 | 1.81E-15 | 26034342785 |
| PNN Num/Area | Exposure | 0.209 | 0.648 | 0.002 | 1.93E-15 | 1406470604 |
|  | Treatment | 1.709 | 0.191 | <0.001 | 5.16E-22 | 17859.937 |
|  | Sex | 0.603 | 0.438 | <0.001 | 1.49E-18 | 5.14E+07 |
|  | Exp. X Treatment | 1.422 | 0.233 | 20533798064 | 2.31E-07 | 1.83E+27 |
| PNN Size | Exposure | 0.106 | 0.744 | 0.003 | 1.69E-18 | 5.03E+12 |
|  | Treatment | 0.82 | 0.365 | 20579212.04 | 3.02E-09 | 1.40E+23 |
|  | Sex | 2.771 | 0.096 | 1.50E+14 | 0.003 | 7.35E+30 |
|  | Exp. X Treatment | 0.056 | 0.813 | 375.766 | 1.75E-19 | 8.05E+23 |
| Number of PV+ cells surrounded by PNNs | Exposure | 1.639 | 0.201 | <0.001 | 1.42E-09 | 71.871 |
|  | Treatment | 1.805 | 0.179 | <0.001 | 2.22E-10 | 63.303 |
|  | Sex | 0.371 | 0.542 | 0.017 | 3.10E-08 | 8849.61 |
|  | Exp. X Treatment | 3.321 | 0.068 | 11741722.59 | 0.293 | 4.71E+14 |
| PV+ cell size when surrounded by PNNs | Exposure | 0.143 | 0.705 | 84.867 | 8.56E-09 | 8.42E+11 |
|  | Treatment | 0.071 | 0.789 | 0.029 | 1.53E-13 | 5483293213 |
|  | Sex | 0.035 | 0.851 | 10.546 | 2.11E-10 | 5.27E+11 |
|  | Exp. X Treatment | 0.041 | 0.839 | 33.142 | 7.56E-14 | 1.45E+16 |
| Number of PNNs surrounding PV+ cells | Exposure | 1.71 | 0.191 | <0.001 | 2.46E-09 | 52.36 |
|  | Treatment | 1.964 | 0.161 | <0.001 | 3.31E-10 | 37.551 |
|  | Sex | 0.414 | 0.52 | 0.015 | 4.56E-08 | 5172.483 |
|  | Exp. X Treatment | 3.485 | 0.062 | 9722204.851 | 0.448 | 2.11E+14 |
| PNN size when surrounding PV+ cells | Exposure | 1.138 | 0.286 | 2068884590 | 1.58E-08 | 2.71E+26 |
|  | Treatment | 1.502 | 0.22 | 2.86E+11 | 1.37E-07 | 5.98E+29 |
|  | Sex | 2.414 | 0.12 | 2.03E+15 | 9.91E-05 | 4.16E+34 |
|  | Exp. X Treatment | 1.344 | 0.246 | <0.001 | 2.00E-39 | 8623664314 |
| Ratio of PV+ cells surrounded by PNNs (PV+PNN/PV total) | Exposure | 0.29 | 0.59 | 0.081 | 8.75E-06 | 755.02 |
|  | Treatment | 0.485 | 0.486 | 0.031 | 1.75E-06 | 545.825 |
|  | Sex | 0.156 | 0.693 | 0.14 | 7.91E-06 | 2464.34 |
|  | Exp. X Treatment | 1.411 | 0.235 | 2607.886 | 0.006 | 1130660895 |
| Ratio of PNNs that surround PV+ cells (PNN+PV/PNN total) | Exposure | 1.558 | 0.212 | <0.001 | 1.75E-08 | 52.615 |
|  | Treatment | 2.717 | 0.099 | <0.001 | 1.44E-10 | 7.073 |
|  | Sex | 0.239 | 0.625 | 18.441 | 0 | 2173939.447 |
|  | Exp. X Treatment | 7.017 | **0.008** | 2402609185 | 275.395 | 2.10E+16 |
| **c. Infralimbic Cortex** |  |  |  |  |  |  |
| **Variable** | **Effect** | **Wald Chi-square** | ***p*** | **Exp(B)** | **95% Wald Confidence Interval** |  |
|  |  |  |  |  | **Lower** | **Higher** |
| PV+ Num/Area | Exposure | 0.307 | 0.58 | 0.003 | 1.73E-12 | 3786202.686 |
|  | Treatment | 0.067 | 0.796 | 0.043 | 1.96E-12 | 959765156 |
|  | Sex | 0.248 | 0.618 | 312.637 | 4.80E-08 | 2.04E+12 |
|  | Exp. X Treatment | 2.329 | 0.127 | 28724448979 | 0.001 | 7.76E+23 |
| PV+ Cell Size | Exposure | 1.862 | 0.172 | 20180115.29 | 0.001 | 6.27E+17 |
|  | Treatment | 3.459 | 0.063 | 45293581923 | 0.267 | 7.68E+21 |
|  | Sex | 0.44 | 0.507 | 6324.325 | 3.73E-08 | 1.07E+15 |
|  | Exp. X Treatment | 5.979 | **0.014** | <0.001 | 2.88E-35 | 0 |
| PNN Num/Area | Exposure | 0 | 0.988 | 0.812 | 3.51E-12 | 1.88E+11 |
|  | Treatment | 0.334 | 0.563 | <0.001 | 5.11E-16 | 213632645.3 |
|  | Sex | 0.002 | 0.967 | 0.567 | 8.76E-13 | 3.66E+11 |
|  | Exp. X Treatment | 1.568 | 0.211 | 14614743353 | 1.80E-06 | 1.19E+26 |
| PNN Size | Exposure | 0.067 | 0.796 | 564.423 | 8.46E-19 | 3.77E+23 |
|  | Treatment | 0.342 | 0.558 | <0.001 | 1.15E-29 | 4.30E+15 |
|  | Sex | 0.337 | 0.562 | <0.001 | 1.29E-29 | 4.85E+15 |
|  | Exp. X Treatment | 0.042 | 0.837 | <0.001 | 2.01E-34 | 2.03E+27 |
| Number of PV+ cells surrounded by PNNs | Exposure | 0.159 | 0.69 | 0.043 | 8.51E-09 | 218100.015 |
|  | Treatment | 0.011 | 0.917 | 0.414 | 2.76E-08 | 6201566.674 |
|  | Sex | 0.278 | 0.598 | 85.063 | 5.68E-06 | 1274054350 |
|  | Exp. X Treatment | 2.963 | 0.085 | 231921046.3 | 0.069 | 7.76E+27 |
| PV+ cell size when surrounded by PNNs | Exposure | 1.53 | 0.216 | 218245518.3 | 1.34E-05 | 3.57E+21 |
|  | Treatment | 1.881 | 0.17 | 7867437281 | 5.68E-05 | 1.09E+24 |
|  | Sex | 0.252 | 0.616 | 4157.654 | 3.00E-11 | 5.76E+17 |
|  | Exp. X Treatment | 4.344 | **0.037** | <0.001 | 1.16E-04 | 0.059 |
| Number of PNNs surrounding PV+ cells | Exposure | 0.228 | 0.633 | 0.024 | 5.46E-09 | 106238.095 |
|  | Treatment | 0.011 | 0.915 | 0.409 | 3.17E-08 | 5285787.388 |
|  | Sex | 0.173 | 0.677 | 32.284 | 2.50E-06 | 417063273.2 |
|  | Exp. X Treatment | 2.809 | 0.094 | 118074622.8 | 0.043 | 3.25E+17 |
| PNN size when surrounding PV+ cells | Exposure | 0.087 | 0.768 | 790.488 | 4.01E-17 | 1.56E+22 |
|  | Treatment | 0.057 | 0.812 | 325.085 | 7.27E-19 | 1.45E+23 |
|  | Sex | 0.149 | 0.699 | <0.001 | 1.92E-25 | 3.83E+16 |
|  | Exp. X Treatment | 1.462 | 0.227 | <0.001 | 2.02E-46 | 67361691310 |
| Ratio of PV+ cells surrounded by PNNs (PV+PNN/PV total) | Exposure | 1.528 | 0.216 | 0.004 | 4.57E-07 | 27.264 |
|  | Treatment | 0.258 | 0.612 | 13.025 | 0.001 | 261763.993 |
|  | Sex | 0.063 | 0.802 | 0.298 | 2.29E-05 | 3875.792 |
|  | Exp. X Treatment | 1.653 | 0.199 | 3604.226 | 0.014 | 952828777.2 |
| Ratio of PNNs that surround PV+ cells (PNN+PV/PNN total) | Exposure | 0.429 | 0.513 | 0.002 | 2.23E-11 | 206838.435 |
|  | Treatment | 0.054 | 0.816 | 0.103 | 5.20E-10 | 20389825.06 |
|  | Sex | 0.523 | 0.47 | 1152.336 | 5.82E-06 | 2.28E+11 |
|  | Exp. X Treatment | 0.67 | 0.413 | 64111.081 | 1.96E-07 | 2.09E+16 |

In the ACC (Figure 2), GLM revealed a significant Exposure X Treatment interaction for PV+ cell number (p=0.020). Pairwise comparisons indicated trends toward reduced PV+ cell number following active stimulation in controls (CON+Sham vs. CON+Active, p=0.059; CON+Active vs. AIE+Active, p=0.057; Figure 2B). No significant effects were observed for PV+ cell size (Figure 2C), PNN number or size (Figure 2D and 2E, respectively), or any PV/PNN co-labeling measures (data not shown). Additionally, no significant effects of sex were detected.

**Figure 2.**
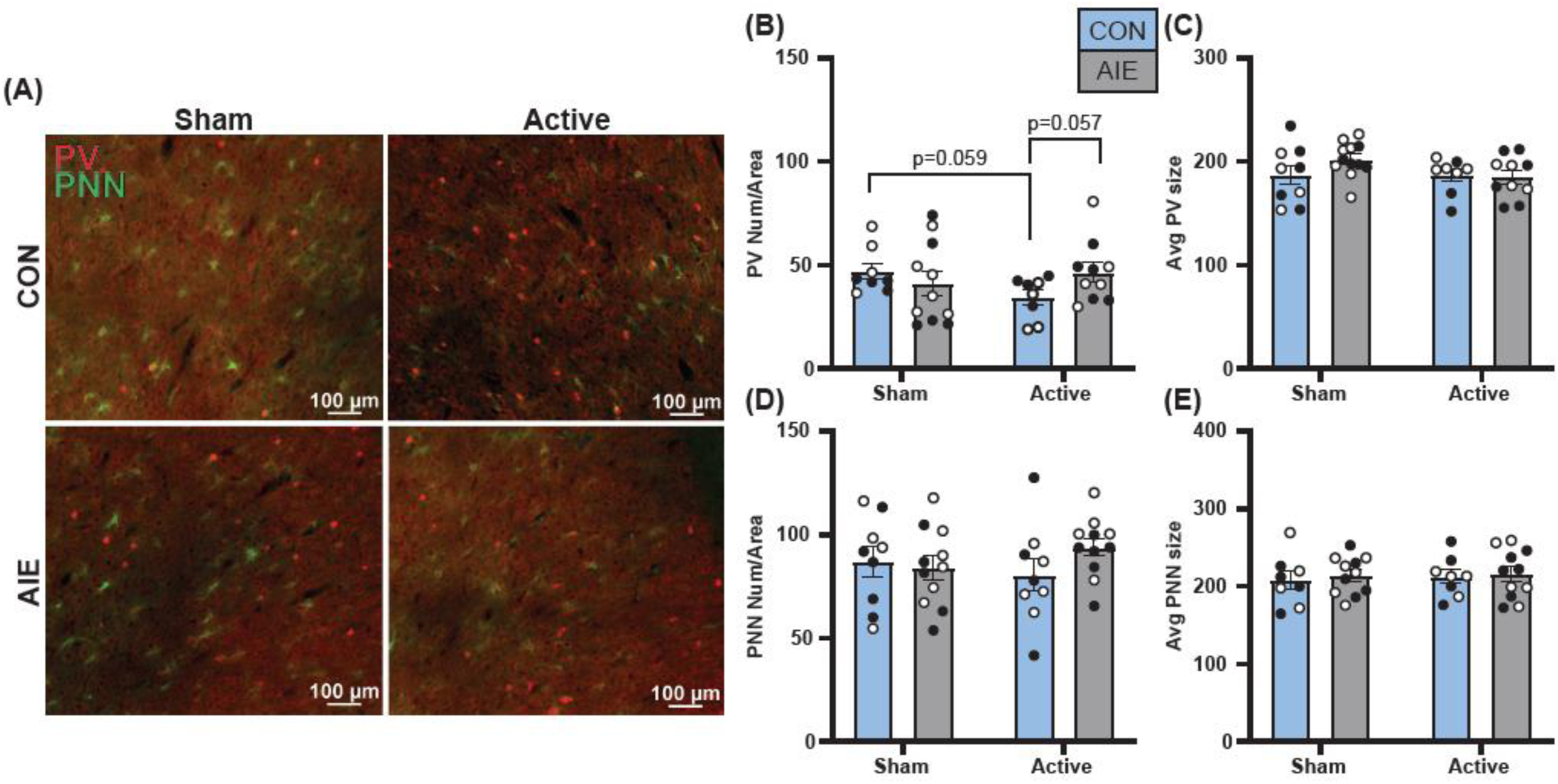
tACS does not significantly impact PV+ cells or PNNs in the anterior cingulate cortex (ACC). (A) Representative image showing immunofluorescence of PV (red) and PNN (green) staining in all four groups. (B) There was a significant Exposure X Treatment interaction for PV+ cell number (p=0.020; cells/mm^2^), and pairwise comparisons indicated trends toward reduced PV+ cell number following active stimulation in controls (CON+Sham vs. CON+Active, p=0.059; CON+Active vs. AIE+Active, p=0.057). There were no group differences in (C) the size of PV+ cells (pixels/number), (D) the number of PNNs (matrices/mm^2^), or (E) the size of PNNs (pixels/number). All data are shown as group mean ± SEM. White circles indicate females; black circles indicate males.

In the PL (Figure 3), GLM revealed a significant Exposure X Treatment interaction for PV+ cell number (p=0.032). AIE exposure reduced PV+ cell number relative to controls (CON+Sham vs. AIE+Sham, p=0.041), while active stimulation normalized PV+ cell number in AIE rats (AIE+Sham vs. AIE+Active, p=0.035; Figure 3B). Trends suggested reduced PV+ cell number following active stimulation in controls (CON+Sham vs. CON+Active, p=0.057; CON+Active vs. AIE+Active, p=0.052). No group differences were observed in PV+ cell size, PNN number, PNN size (Figure 3C-E), any PV/PNN co-labeling measures (data not shown), nor were there any sex effects.

**Figure 3.**
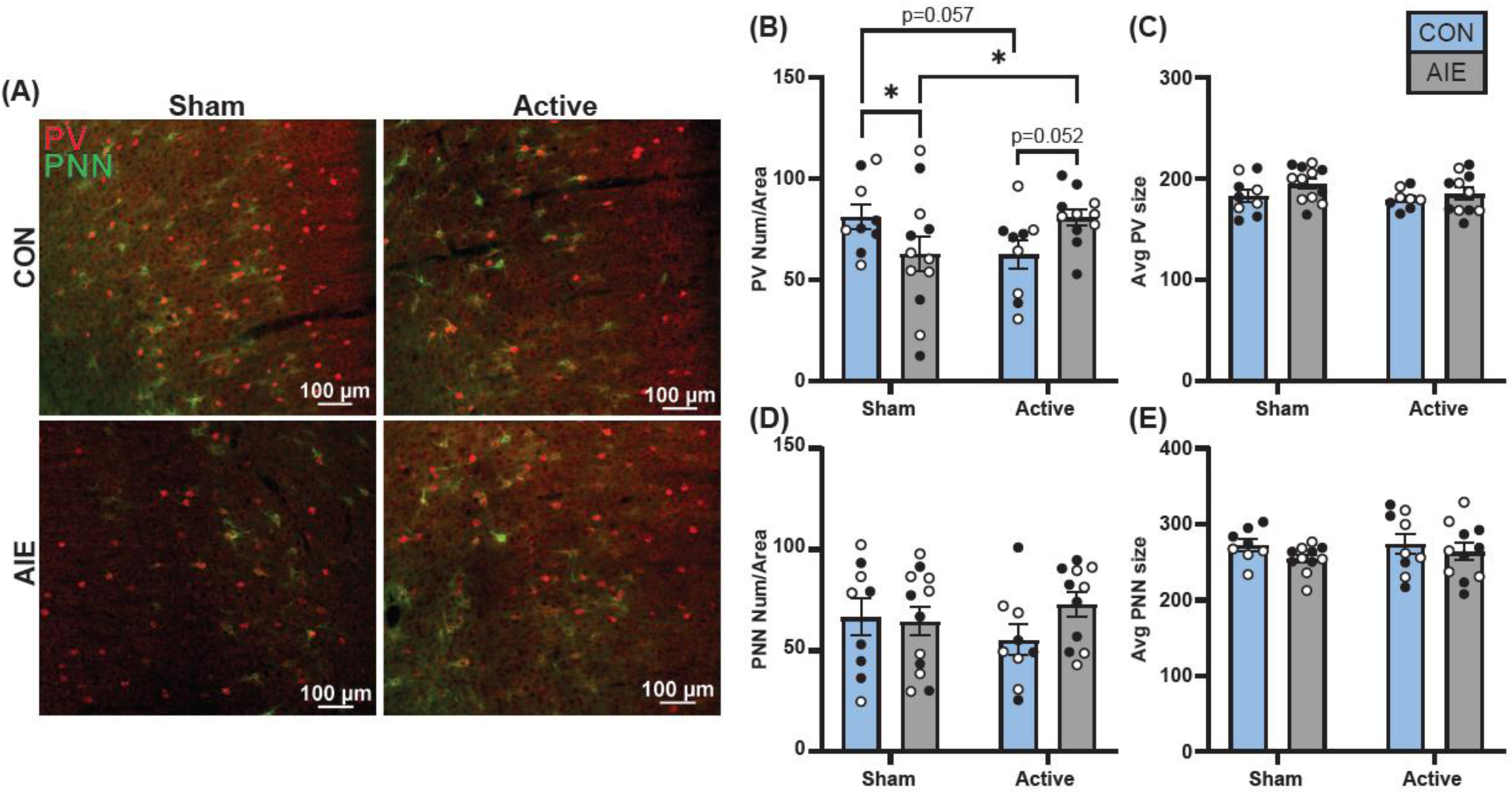
Active tACS ameliorates AIE-induced reductions in PV+ cell number in the prelimbic (PL) cortex. (A) Representative image showing immunofluorescence of PV (red) and PNN (green) staining in all four groups. (B) AIE exposure reduced PV+ cell number (cells/mm^2^) relative to controls (CON+Sham vs. AIE+Sham, p=0.041), while active stimulation normalized PV+ cell number in AIE rats (AIE+Sham vs. AIE+Active, p=0.035). There were no differences between groups in (C) the size of PV+ cells (pixels/number), (D) the number of PNNs (matrices/mm^2^), or (E) the size of PNNs (pixels/number). All data are shown as group mean ± SEM. White circles indicate females; black circles indicate males.

In the IL (Figure 4), neither AIE exposure nor tACS treatment altered PV+ cell number (Figure 4B). However, a significant Exposure X Treatment interaction was detected for PV+ cell size (p=0.014), with active stimulation reducing PV+ cell size in AIE rats compared to both CON+Active (p=0.010) and AIE+Sham rats (p=0.026; Figure 4C). A trend toward increased PV+ cell size was observed in stimulated controls (CON+Sham vs. CON+Active, p=0.052). A similar relationship emerged when isolating the PV+ cells enclosed by PNNs (p=0.037), with active stimulation reducing PV+ cell size in AIE rats compared to AIE+Sham animals (p=0.034; data not shown). No effects were observed for PNN number or size (Figure 4D and 4E, respectively) or for other PV/PNN co-labeling measures (data not shown), and there were no effects of sex.

**Figure 4.**
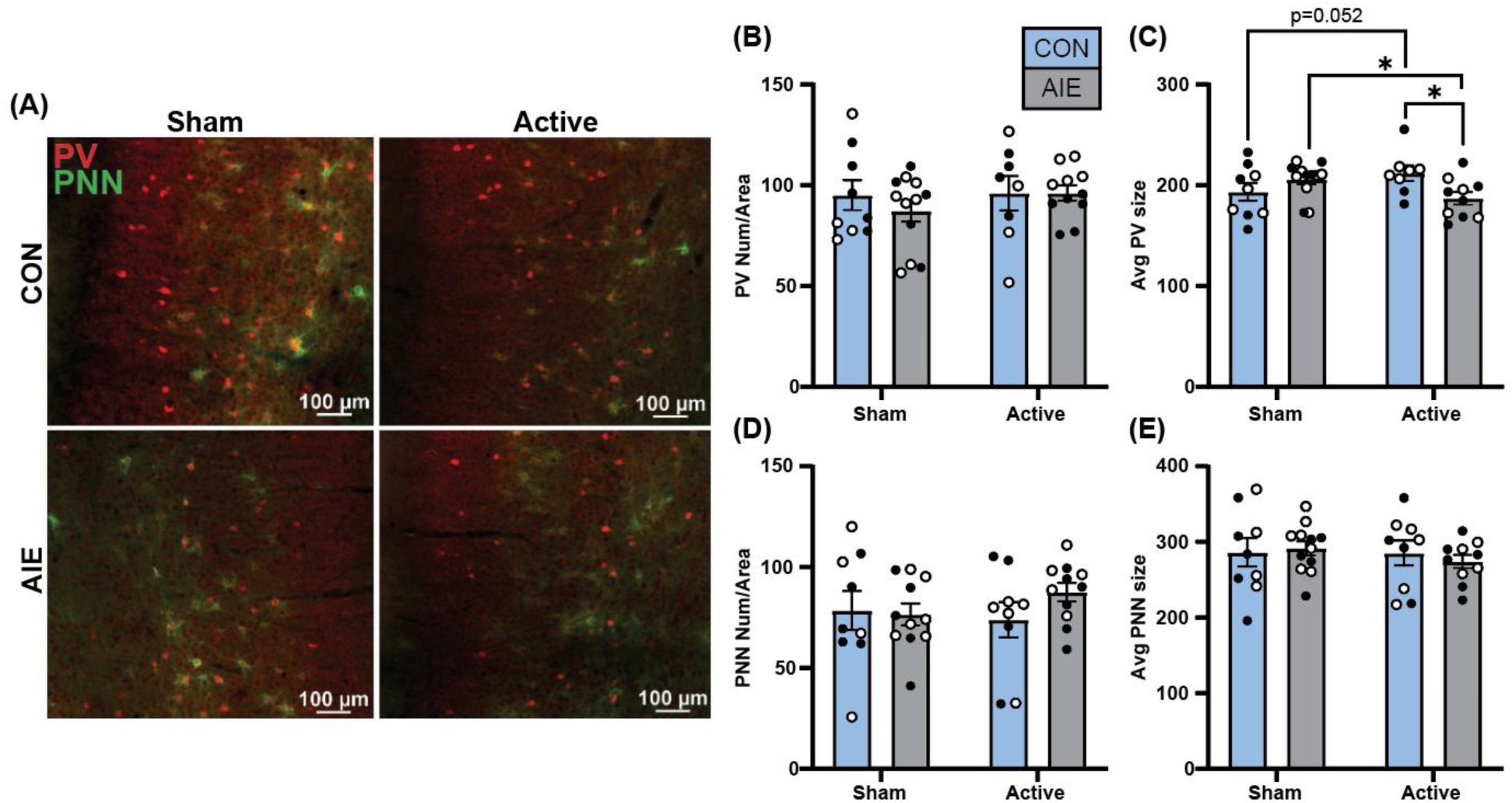
Active tACS reduces PV+ cell size in AIE-exposed rats in the infralimbic (IL) cortex. (A) Representative image showing immunofluorescence of PV (red) and PNN (green) staining in all four groups. (B) There were no group differences in PV+ cell number (cells/mm^2^). (C) Active stimulation reduced PV+ cell size (pixels/number) in AIE rats compared to both CON+Active (p=0.010) and AIE+Sham rats (p=0.026). There were no differences between groups in (D) the number of PNNs (matrices/mm^2^) or (E) the size of PNNs (pixels/number). All data are shown as group mean ± SEM. White circles indicate females; black circles indicate males.

### Effects of AIE and tACS on recognition memory

We assessed the impact of tACS on memory both acutely (same day) and persistently (one week later) in separate groups of rats (see Table 2 for full statistics). At the acute timepoint (Figure 5A), two-way ANOVA revealed significant main effects of Exposure (F_1,42_ = 9.465, p=0.004) and Treatment (F_1,42_ = 10.709, p=0.002) for percent preference of the novel object, indicating that AIE exposure impaired NOR memory compared to control exposure, and that active stimulation decreased novel object preference compared to sham stimulation. At the one-week post stimulation timepoint (Figure 5B), two-way ANOVA revealed a significant Exposure X Treatment interaction (F_1,28_ = 10.820, p=0.003) for percent preference. Tukey’s multiple comparisons tests indicated that AIE impaired NOR memory (AIE+Sham vs. CON+Sham, p=0.013; AIE+Sham vs. CON+Active; p=0.002). Notably, active stimulation reversed the AIE-induced deficit in NOR, with AIE+Active rats performing significantly better than AIE+Sham rats (p<0.0001). At both timepoints, there were no differences between males and females.

**Figure 5.**
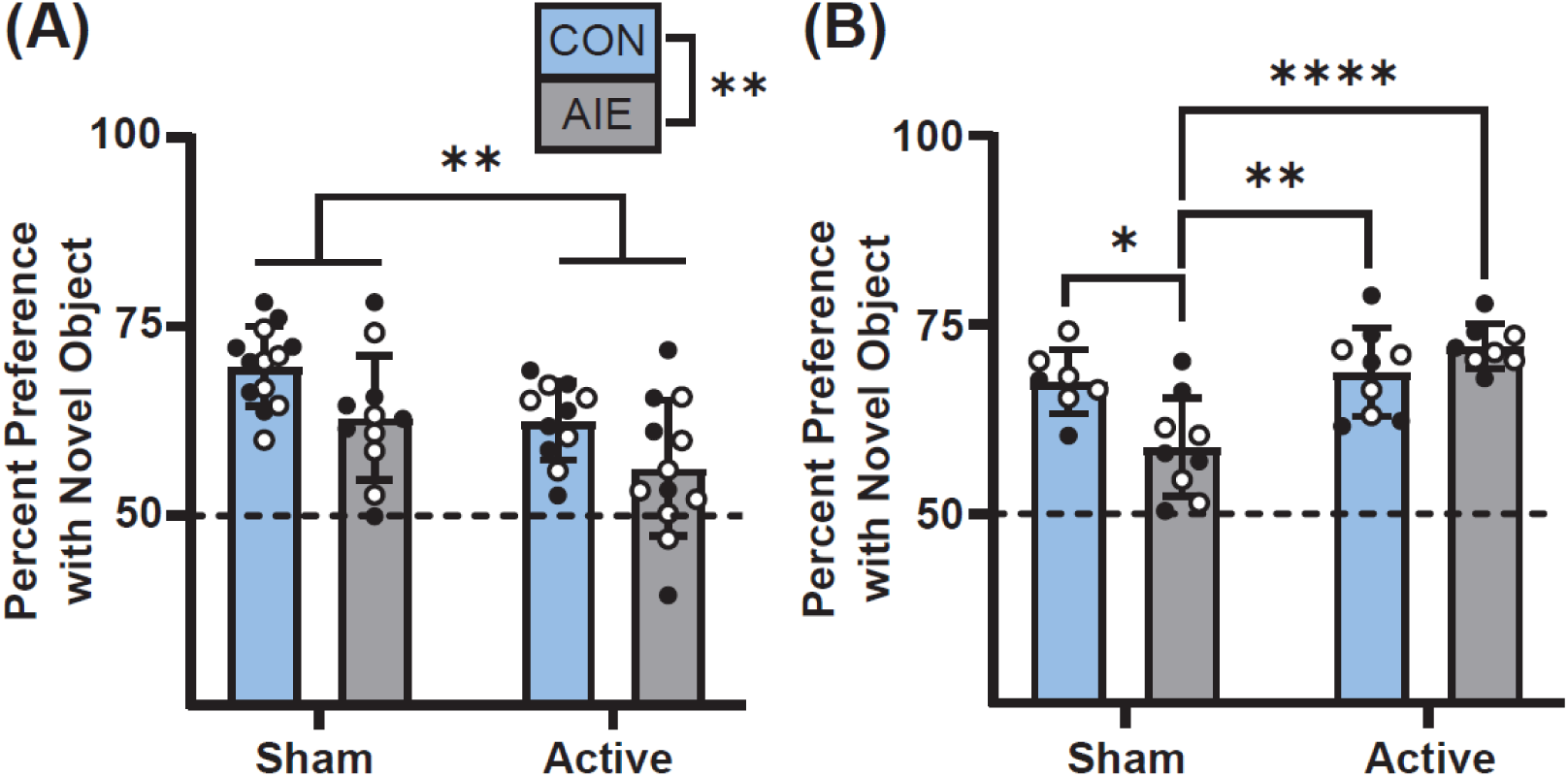
tACS differentially impacts recognition memory at acute and persistent timepoints. (A) At the acute timepoint, there were main effects of Exposure (p=0.004) and Treatment (p=0.002) where AIE rats performed worse than their CON counterparts, and Active tACS reduced percent preference for the novel object compared to Sham stimulation. (B) One week post-stimulation, there was a significant Exposure X Treatment interaction (p=0.003), where AIE+Sham rats showed lower percent preference than both CON+Sham (p=0.013) and CON+Active (p=0.002) rats. Active tACS reversed the AIE-induced deficit in preference, with AIE+Active rats showing higher preference for the novel object than AIE+Sham rats (p<0.0001). All data are shown as group mean ± SEM. White circles indicate females; black circles indicate males.

**Table 2.** All behavioral results. Two-way analyses of variance were used with Exposure (AIE, CON) and Treatment (Active, Sham) as between-subjects factors and Sex (Male, Female) as a covariate. Significant p-values (p < 0.05) are bolded and red.

| **Behavior** |  |  |  |  |  |
| **a. Novel Object Recognition** |  |  |  |  |  |
| **Variable** | **Effect** | **Mean Square** | **F** | ***p*** | **Observed Power<sup>b</sup>** |
|---|---|---|---|---|---|
| Percent Preference: Acute Timepoint | Exposure | 475.116 | 9.465 | **0.004** | 0.852 |
|  | Treatment | 537.527 | 10.709 | **0.002** | 0.892 |
|  | Sex | 49.77 | 0.992 | 0.325 | 0.164 |
|  | Exp. X Treatment | 2.206 | 0.044 | 0.835 | 0.055 |
| Percent Preference: Persistent Timepoint | Exposure | 58.353 | 2.098 | 0.159 | 0.288 |
|  | Treatment | 419.066 | 15.068 | **<0.001** | 0.963 |
|  | Sex | 5.214 | 0.187 | 0.668 | 0.07 |
|  | Exp. X Treatment | 300.927 | 10.82 | **0.003** | 0.888 |
| **b. Open Field Behaviors** |  |  |  |  |  |
| **Variable** | **Effect** | **Mean Square** | **F** | ***p*** | **Observed Power<sup>b</sup>** |
| Latency to enter center: Acute Timepoint | Exposure | 5.215 | 0.013 | 0.911 | 0.051 |
|  | Treatment | 3069.388 | 7.463 | **0.009** | 0.762 |
|  | Sex | 943.573 | 2.294 | 0.137 | 0.316 |
|  | Exp. X Treatment | 83.237 | 0.202 | 0.655 | 0.072 |
| Total Duration in Center: Acute Timepoint | Exposure | 31.299 | 0.046 | 0.83 | 0.055 |
|  | Treatment | 3130.64 | 4.642 | **0.036** | 0.559 |
|  | Sex | 1221.168 | 1.811 | 0.185 | 0.261 |
|  | Exp. X Treatment | 1302.396 | 1.931 | 0.171 | 0.275 |
| Bin 1 Duration in Center: Acute Timepoint | Exposure | 117.286 | 1.752 | 0.192 | 0.254 |
|  | Treatment | 793.811 | 11.858 | **0.001** | 0.921 |
|  | Sex | 134.705 | 2.012 | 0.163 | 0.284 |
|  | Exp. X Treatment | 23.253 | 0.347 | 0.559 | 0.089 |
| Total Line Crossings: Acute Timepoint | Exposure | 1573.657 | 0.586 | 0.448 | 0.116 |
|  | Treatment | 14.593 | 0.005 | 0.942 | 0.051 |
|  | Sex | 60869.018 | 22.657 | **<0.001** | 0.997 |
|  | Exp. X Treatment | 1234.472 | 0.429 | 0.501 | 0.102 |
| Latency to enter center: Persistent Timepoint | Exposure | 0.853 | 0.002 | 0.964 | 0.05 |
|  | Treatment | 81.681 | 0.195 | 0.662 | 0.071 |
|  | Sex | 55.945 | 0.133 | 0.717 | 0.064 |
|  | Exp. X Treatment | 276.975 | 0.66 | 0.423 | 0.124 |
| Total Duration in Center: Persistent Timepoint | Exposure | 155.987 | 0.207 | 0.652 | 0.073 |
|  | Treatment | 1157.202 | 1.536 | 0.224 | 0.225 |
|  | Sex | 1433.967 | 1.903 | 0.177 | 0.268 |
|  | Exp. X Treatment | 14.018 | 0.019 | 0.892 | 0.052 |
| Bin 1 Duration in Center: Persistent Timepoint | Exposure | 11.869 | 0.235 | 0.631 | 0.076 |
|  | Treatment | 239.83 | 4.757 | **0.037** | 0.561 |
|  | Sex | 14.147 | 0.281 | 0.6 | 0.081 |
|  | Exp. X Treatment | 7.29 | 0.145 | 0.706 | 0.066 |
| Total Line Crossings: Persistent Timepoint | Exposure | 12892.992 | 6.56 | **0.016** | 0.697 |
|  | Treatment | 1060.787 | 0.54 | 0.468 | 0.11 |
|  | Sex | 4458.031 | 2.268 | 0.143 | 0.308 |
|  | Exp. X Treatment | 1599.42 | 0.814 | 0.374 | 0.141 |

### Effects of AIE and tACS on anxiety-like behaviors

Using spontaneous locomotion in the arena on day 1 of the NOR test, we assessed the acute (same day) and persistent (one week later) impact of tACS on anxiety-like behavior in separate groups of rats (full statistics in Table 2). At the acute timepoint, two-way ANOVA revealed that a single session of active tACS reduced the latency to enter the center of the open-field arena compared with sham stimulation (F_1,44_ = 7.463, p=0.009; Figure 6A). Active stimulation also increased time spent in the center, both across the entire session (F_1,46_ = 4.642, p=0.036) and during the first 2.5-minute time bin (F_1,45_ = 11.858, p=0.001; Figure 6B-C). Importantly, total line crossings were not affected by Treatment or Exposure; however, females exhibited greater locomotor activity than males, as indicated by increased line crossings (F_1,46_ = 22.657, p<0.001; Figure 6D).

**Figure 6.**
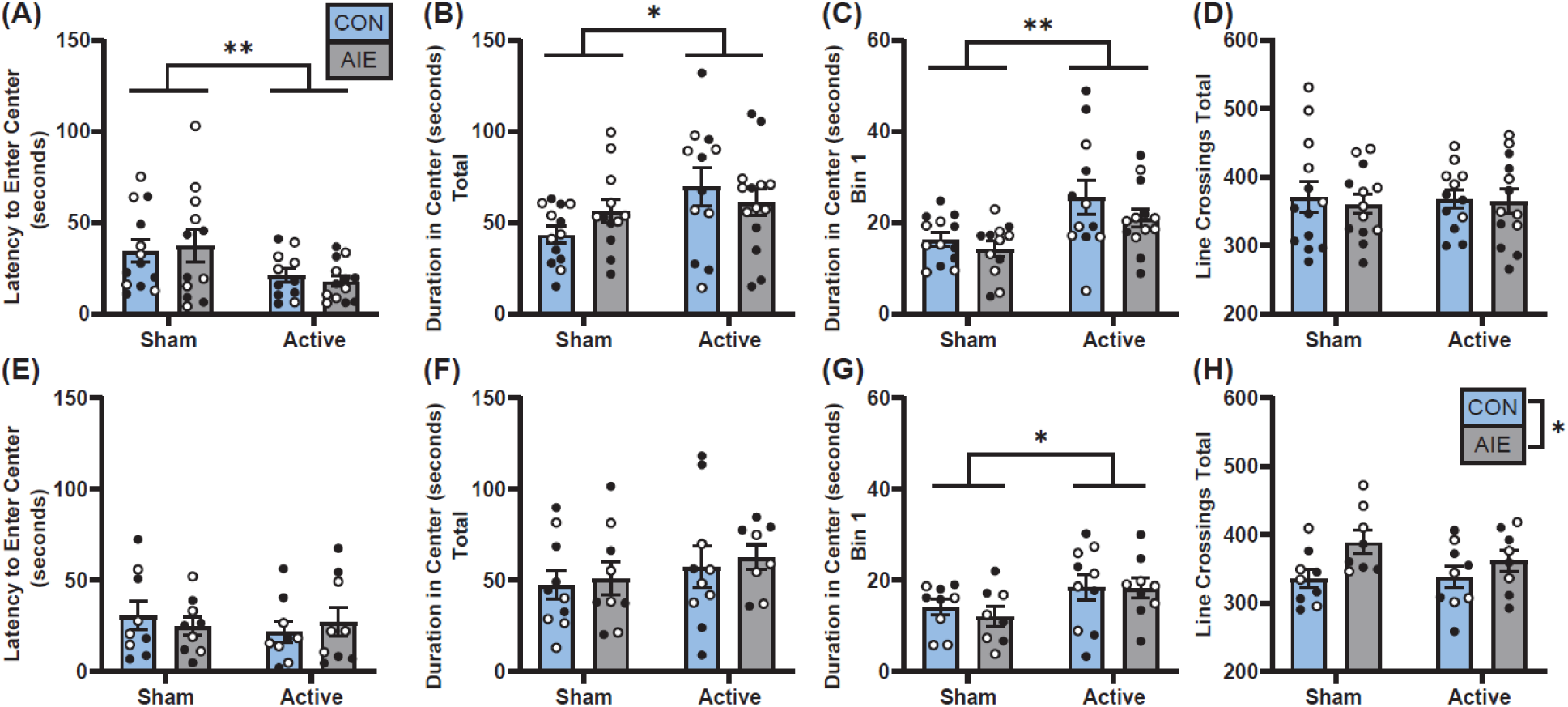
tACS reduces anxiety-like behaviors at acute and persistent timepoints. At the acute timepoint, a single session of active tACS (A) reduced the latency to enter the center of the open-field arena compared with sham stimulation (p=0.009), (B) increased the total duration of time in the center of the arena compared to sham stimulation (p=0.036), and (C) increased the time in the center in the first 2.5-minute time bin (p=0.001). (D) There were no effects of Exposure or Treatment in the total number of line crossings following a single session of stimulation. There was a main effect of sex where females made more line crossings than males (p<0.001). One week post-stimulation, there were no differences among groups (E) in the latency to enter the center of the arena or (F) the total duration of time in the center of the arena. (G) During the first 2.5-minute time bin, active stimulation led to increased time spent in the center compared with sham stimulation (p=0.037). (H) There were no effects of Sex or Treatment in the total number of line crossings, however AIE rats had more line crossings than CON rats (p=0.016). All data are shown as group mean ± SEM. White circles indicate females; black circles indicate males.

At one-week post-stimulation, no group differences were observed in latency to enter the center or in total time spent in the center of the arena (Figure 6E, 6F). However, rats that received active stimulation spent more time in the center during the first 2.5-minute time bin compared with sham-treated rats (F_1,31_ = 4.757, p=0.037; Figure 6G). Total line crossings were not affected by Treatment or Sex, but a significant main effect of Exposure (F_1,29_ = 6.560, p=0.016) showed that AIE exposed rats exhibited greater locomotor activity than control rats (Figure 6H).

## DISCUSSION

In the present study, we found that mPFC-targeted, gamma-frequency tACS mitigated AIE-induced alterations in PV+ interneurons without impacting PNNs. We also found that AIE exposure reduced recognition memory of a novel object, and that gamma tACS ameliorated this deficit when assessed one week after stimulation. Furthermore, active stimulation decreased anxiety-like behaviors in the open-field test.

We investigated PV+ interneurons as a primary target of tACS because they are essential for generating gamma oscillations (13–15) and are highly vulnerable to adolescent alcohol exposure. Multiple studies have shown that AIE reduces PV+ cell number, expression, or cell size in several brain regions, including the mPFC, with some effects persisting into adulthood (21,41–44). In addition, AIE impairs PV+ interneuron function by reducing intrinsic excitability and altering excitatory/inhibitory balance at basolateral amygdala inputs onto mPFC PV+ cells (45). Although findings vary across studies, likely due to methodological factors such as tissue collection timing and regional or sex differences (21,40,46,47), the collective literature demonstrates that PV+ interneurons are vulnerable to alcohol-induced modifications.

We observed region-specific effects of AIE exposure and tACS treatment on PV+ interneuron metrics. In the PL cortex, AIE exposure reduced the number of PV+ cells, and active stimulation ameliorated these reductions. In contrast, in the IL cortex, active stimulation decreased PV+ cell size in AIE-exposed rats without affecting cell number. We also observed trends toward reduced numbers of PV+ cells in control animals receiving active stimulation in both the PL and ACC cortices. Notably, significant stimulation-related effects were detected in the PL and IL cortices rather than in the ACC, despite its closer proximity to the stimulation site. These findings suggest that the histological effects of gamma-frequency tACS may depend on stimulation-induced network dynamics or an optimal distance from the stimulation source. Although neuronal function was not assessed in the present study, it remains possible that tACS-induced changes in oscillatory activity can occur in the absence of detectable histological alterations.

Consistent with this possibility, previous work using the same stimulation parameters demonstrated functional changes measured by local field potential recordings, where cocaine exposure reduced coherence between the PL cortex and nucleus accumbens, a deficit that was restored following mPFC-targeted gamma-frequency tACS (37). Future studies could address this gap by assessing markers of neuronal activity, such as immediate early genes, to determine whether tACS alters neural activation patterns in the absence of structural changes. Such approaches could also extend to downstream brain regions to better characterize network-wide effects of mPFC-targeted stimulation.

Although the mechanisms underlying AIE- and tACS-induced changes in PV+ cell numbers and size remain unclear, they may involve epigenetic regulation. AIE exposure has been shown to alter epigenetic processes, including histone methylation and acetylation, which regulate gene expression (11). For example, AIE exposure increases histone deacetylase activity, reduces histone H3 acetylation in the hippocampus (48), and increases the gene-silencing marker demethylated histone H3K9 (49). In the PL cortex, AIE exposure also alters transcriptional profiles, increasing mRNA expression of several neuroinflammatory markers (50). Additionally, AIE-induced repression of basal forebrain cholinergic neuron phenotypes can be reversed by exercise (51,52) and anti-inflammatory or epigenetic interventions (53–55). Thus, if the AIE-induced reduction in PV+ cell numbers and size observed in the PL and IL cortices reflects epigenetic suppression rather than cell loss, then tACS may restore PV expression through epigenetic mechanisms. However, whether the AIE- and tACS-induced changes are transient, and the functional relevance of these histological changes remain unclear.

In the present study, we found no changes in mPFC PNNs. This finding contrasts with previous reports by us and others that AIE exposure increased the number of PNNs in the mPFC (40,56,57), although this effect varied by subregion and cortical layer (57). PNNs are highly dynamic structures that are influenced by a variety of factors including stress (58), memory retrieval (59), drug exposure (40,60,61), behavioral training (21), and even diurnal fluctuations (47). As such, when we assessed the mPFC from male and female rats after extensive behavioral testing, we no longer observed AIE-induced changes in PNNs in the mPFC, although we observed increased PNN numbers and a greater proportion of PV+ neurons ensheathed by PNNs in the anterior insular cortex in the same AIE-exposed rats (21). Although the animals in the current study did not receive formal behavioral training, they underwent surgery and extensive handling associated with tACS procedures. Thus, the absence of AIE-induced PNN alterations in the mPFC in the current study may have constrained the ability to detect tACS-induced effects.

Consistent with previous findings (9,10,62), AIE exposure produced persistent recognition memory deficits in the NOR task. Importantly, we found that the effects of tACS depended on the timing of the behavioral assessment. While tACS impaired same-day recognition memory performance, when assessed one week after stimulation, tACS attenuated the AIE-induced deficit in object recognition memory. The acute impairment is not entirely unexpected. Clinically, tACS administered during the Hidden Association Between Images Task (HABIT), a measure of stimulus-response learning and habit formation, impaired performance in healthy control participants (29), while improving performance in subjects with a history of substance use disorder. Similarly, in a preclinical study, tACS weakened Pavlovian conditioning in drug-naïve rats but restored behavioral deficits in drug-exposed animals (37). Together, these findings suggest that acute stimulation may transiently disrupt neural processing, potentially through overstimulation of already engaged circuits. In contrast, delayed assessment following tACS revealed improved recognition memory in AIE-exposed animals without affecting performance in control animals. One explanation is that the post-stimulation interval allowed neural circuits to adapt to stimulation-induced changes, thereby normalizing circuit function disrupted by AIE exposure. On the other hand, tACS would disrupt the presumably optimal state of control animals, but this disruption would have resolved a week later, resulting in normal performance.

To interrogate how tACS stimulation was perceived by the animal, we assessed open-field behavior and found that tACS reduced anxiety-like behavior at both acute and persistent timepoints. The effect was strongest immediately following stimulation (acute) and, notably, was observed after only a single tACS session. Although attenuated, an anxiolytic-like effect remained evident one week later, with actively stimulated animals spending more time in the center during the first time bin. Importantly, tACS did not alter overall locomotor activity, indicating that the reduced latency to enter the center and increased center time were not due to increased movement. One consideration is that the NOR task was conducted under low-light conditions to promote object exploration. As the reduced lighting likely decreased the aversive nature of the open field, the behavioral conditions used here may not have been optimal for detecting anxiety-related phenotypes. This could contribute to differences between the current findings and previous reports of increased anxiety-like behavior following AIE exposure (5,63,64), which we did not observe. Future studies evaluating the effects of tACS on anxiety-like behavior should employ paradigms specifically designed to measure anxiety, such as the elevated plus maze or light-dark box.

In conclusion, these findings demonstrate that mPFC-targeted gamma tACS can partially reverse persistent neurobiological and behavioral alterations induced by adolescent alcohol exposure. These results provide the initial foundation for understanding the physiological impact of a promising noninvasive therapeutic method. Future studies integrating behavioral assessments with targeted manipulations will be important for determining whether changes in PV+ interneurons mechanistically contribute to the behavioral effects of AIE exposure and their restoration by tACS.

